# Hidden origami in *Trypanosoma cruzi* nuclei highlights its nonrandom 3D genomic organization

**DOI:** 10.1101/2024.07.01.601582

**Authors:** Natália Karla Bellini, Pedro Leonardo Carvalho de Lima, David da Silva Pires, Julia Pinheiro Chagas da Cunha

## Abstract

The protozoan *Trypanosoma cruzi*, the causative agent of Chagas disease, exhibits polycistronic transcription and unidimensional genome compartmentalization of *core* (conserved) and *disruptive* (virulence factors from multigenic families) genes. Approximately 50% of its genome is repetitive, mainly virulence factor genes. Genomic sequences, including repeats, motifs of architectural proteins, and noncoding RNA loci are crucial for genome folding. Here, we evaluated the genomic features associated with higher-order chromatin organization in *T. cruzi* through extensive computational processing of high-throughput chromosome conformation capture (Hi-C) data, accounting for repetitive regions and improvements in genome annotation. Our study revealed that repetitive DNA (multimapped reads) influences 3D chromatin folding, particularly in determining the boundaries of topologically associated domains (TAD)-like structures. Virulence factor genes, unlike *core* genes, form shorter and more compact TAD-like structures enriched in loops, suggesting a gene expression regulatory mechanism. We found nonprotein-coding RNA loci (e.g., tRNAs) and transcription termination sites preferentially located at the boundaries of the TAD-like structures, while pseudogenes and multigenic family genes located in unstructured genomic regions. Our data indicate 3D clustering of tRNA loci, likely optimizing transcription by RNA polymerase III, and a complex interaction between spliced-leader RNA and 18S rRNA loci. Our findings provide insights into 3D genome organization in *T. cruzi*, contributing to the understanding of supranucleosome-level chromatin organization and suggesting possible links between 3D architecture and gene expression. We draw an analogy to the art of origami (e.g., papers folded into various shapes) resembling the DNA packed in chromatin fibers assuming distinct folds within the nucleus.

**Importance:** Despite the knowledge about the linear genome sequence and the identification of numerous virulence factors in the protozoan parasite *Trypanosoma cruzi*, there has been a limited understanding of how these genomic features are spatially organized within the nucleus and how this organization impacts gene regulation and pathogenicity. By providing a detailed analysis of the three-dimensional chromatin architecture in *T. cruzi*, our study contributed to filling this gap. We deciphered part of the origami structure hidden in the *T. cruzi* nucleus, showing the unidimensional genomic features are nonrandomly organized in the nuclear 3D landscape. We revealed the possible role of non-protein-coding RNA loci (e.g., tRNAs, SL-RNA, and 18S RNA) in shaping the genomic architecture. These findings provide insights into an additional epigenetic layer that may influence gene expression.

**Graphical abstract:** The spatial organization of chromatin within the nuclei of *T. cruzi* and its resemblance to origami art. A. Identification of the 3D nuclear architectures within *T. cruzi* nuclei: topologically associating domains (TADs) and their boundaries; chromatin loops; and 3D networks. Inter- and intrachromosomal interactions reflect DNA‒DNA contacts on the same (*cis*) and between different (*trans*) chromosomes. B. Resemblance between origami art and chromatin folding. Steps “a” to “l” show the process of folding a flat piece of paper from its unidimensional view up to its 3D boat form.

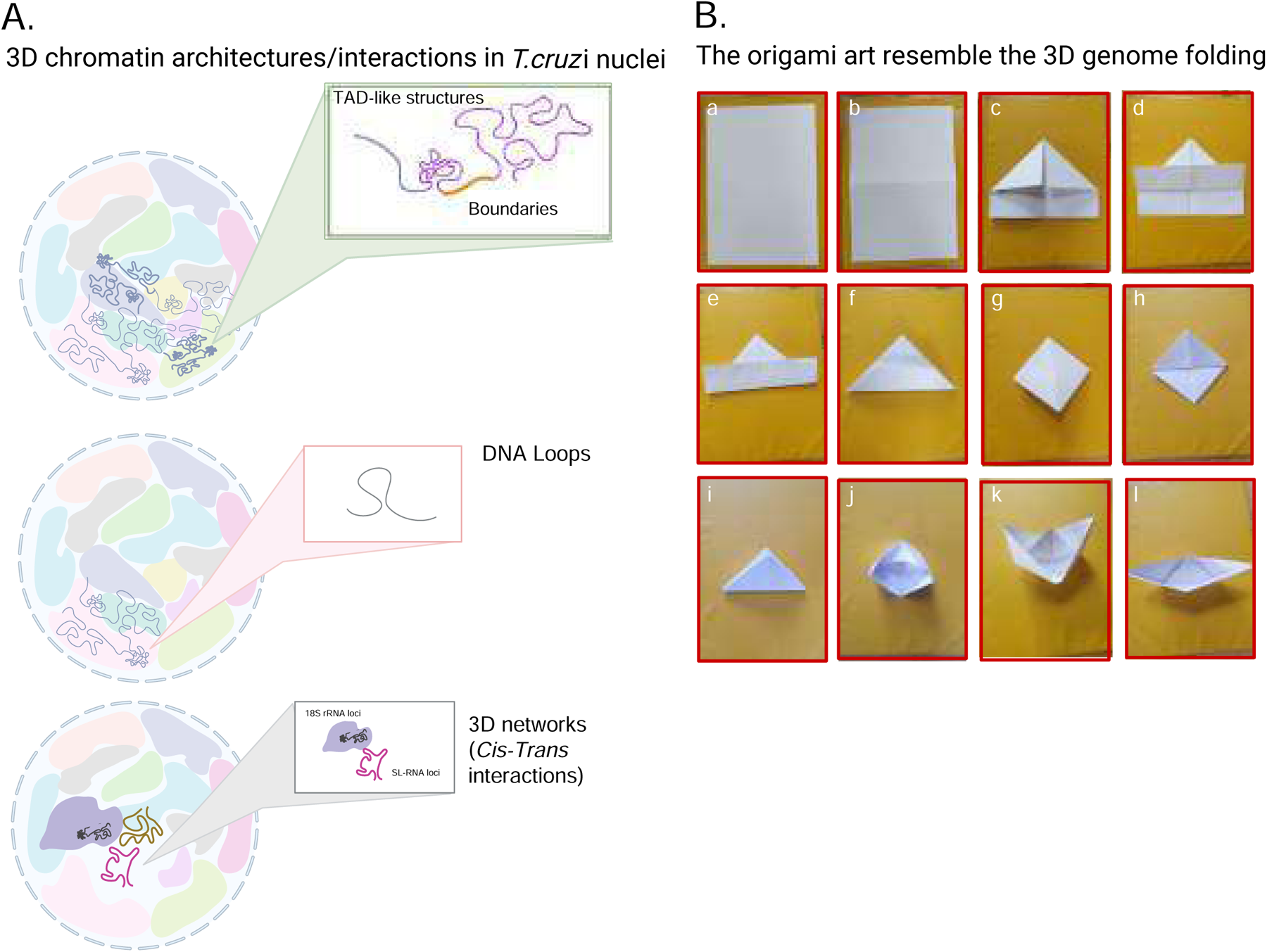

## INTRODUCTION

*Trypanosoma cruzi* is a heteroxenic protozoan parasite and the etiological agent of Chagas disease, which primarily affects Latin American countries. However it has now reached a global distribution, impacting nonendemic regions as well (1). Genomic peculiarities of *T. cruzi* include directional clusters of genes transcribed polycistronically by RNA polymerase II without well-known promoter regions (except for spliced leader (SL)-genes) (2); the absence of introns (except for the poly-A polymerase gene) (3); RNA processing mediated by the trans-splicing machinery (4); the compartmentalization of genes into *core* (conserved, syntenic genes) and *disruptive* (multigene families, nonsyntenic genes); and an abundance of repetitive DNA (5–7). Moreover, a family of genes encoding the virulence factor genes Gp63, DGF and RHS are distributed throughout the genome, positioned within *core* and *disruptive* compartments (7). These gene families are collectively referred to as the *GpDR* group.

Protein production in *T. cruzi* can be controlled at multiple levels. Unlike most eukaryotes, which primarily regulate gene expression at the initiation of transcription, trypanosomatids rely predominantly on posttranscriptional mechanisms, including transcript processing, stability, translation (4), and codon usage bias (8, 9). Additionally, noncoding RNAs (ncRNAs) play significant roles in the regulation of gene expression, contributing to posttranscriptional control mechanisms. As in other eukaryotes, small nuclear RNAs (snRNAs) are essential for mRNA maturation and include U-rich snRNAs that form the spliceosome and SL-RNAs that provide the 39-nt sequence for trans-splicing (10). Ribosomal RNAs (rRNAs) and their genes undergo extensive processing and chemical modifications, which are crucial for their function in the ribosome (11). In addition, both SL-RNA loci and rRNA loci have been associated with 3D nuclear compartmentalization in trypanosomatids and in mammals, which also impacts gene expression (12–14).

Epigenetic factors such as nucleosome occupancy and dynamics (15), open chromatin status (16), DNA modifications such as base J deposition at the TTS (17, 18), histone variant deposition (19, 20) and histone posttranslational modifications have also been demonstrated to be additional players in gene regulation (21).

In *T. cruzi*, epigenetic mechanisms during the differentiation from replicative to nonreplicative forms have been extensively studied. The infective, nonproliferative form shows lower global transcriptional levels of RNA polymerase II associated with nuclear chromatin alterations (22). The chromatin structure is dynamically modulated in its different life forms, with open chromatin in epimastigotes enriched in *core* genes and closed chromatin associated with *disruptive* genes, the virulence factor proteins (16). Histone variants such as H2B.V are found at transcription start sites (TSSs) and often colocalize with tRNA loci (19), while H4.V is enriched at transcription termination sites (TTS) and telomeric regions (Rosón et al. 2024, in preparation). During differentiation, trypomastigotes exhibit increased heterochromatin accompanied by the concentration of euchromatin, mostly in the inner nucleus, and increased nucleosome density at TSS regions, which is correlated with decreased global transcription levels (15, 16).

Open chromatin is essential for gene expression in many organisms (23). Three-dimensional (3D) nuclear architectures, such as active and inactive compartments, are linked to gene expression control (24, 25) and to DNA replication and repair (26). Chromosome conformation capture (3C), introduced by Dekker et al. in 2002 (27), enabled the study of 3D DNA‒DNA interactions within the nucleus. This technique involves cross-linking DNA, digesting it with an endonuclease, and ligating it to preserve 3D contacts, followed by PCR to detect interactions. The method, adapted from nuclear ligation assays by Cullen et al. in 1993 (28), was further advanced by Lieberman-Aiden et al. in 2009 to become the Hi-C method, which uses next-generation sequencing to map genome-wide DNA interactions (24).

In trypanosomatids, the Hi-C technique was first utilized by Muller L. et al. to reveal the 3D nuclear organization of core and subtelomeric regions and centromeres (29). A subsequent study revealed spatial proximity between active variant surface glycoprotein (VSG) expression sites and spliced leader (SL) RNA loci, suggesting that switching among VSG expression sites is linked with their proximity to the SL array in *T. brucei* (12). In 2021, Hi-C was employed for the first time in a *T. cruzi* strain (Brazil clone A4) to aid genome assembly by resolving gaps in short-read and long-read sequences (30). A study linking 3D chromatin structure in *T. cruzi* with DNA methylation, nucleosome positioning, and transcription levels, highlighting the organization of *core* and *disruptive* compartments, was recently published; however, it did not address the impact of repetitive DNA on the 3D nuclear architecture (31).

Drawing an analogy to origami, where paper folds into a variety of shapes, the three-dimensional chromatin organization within cell nuclei similarly adopts diverse and intricate structures. Linear DNA is primarily packed in chromatin fibers, which also assume distinct folds inside the nucleus (32). The two most well-characterized 3D chromatin architectures comprise topologically associated domains (TADs) and chromatin loops (**refer to the graphical abstract**). In the context of 3D origami, folding a flat material brings certain regions to the surface, making them accessible for decoration, while other regions become hidden and less accessible. DNA packaged in chromatin similarly folds into 3D arrangements, bringing distant loci into proximity, while others remain far apart (32). This concept parallels how regions of DNA that are distant in the linear genome rarely interact in the 3D nuclear space, whereas loci that are close in the linear sequence frequently contact each other when folded into a 3D structure. The closer two DNA loci are along the chromosome, the greater their frequency of 3D contact (33). DNA regions that are considerably distant in a one-dimensional (1D) context commonly provide significant cell biology benefits when interacting in a 3D space. Examples of these benefits include the well-characterized enhancers, which facilitate gene regulation by looping to their target promoters despite being far apart in the linear genome (34). Additionally, various protein‒DNA interactions contribute to shaping the chromatin architecture, as seen in the folding patterns influenced by transcription factors and chromatin remodelers (35). These interactions, along with the chromatin compaction landscapes (36), are crucial for the regulation of gene expression and the maintenance of genome stability, illustrating the functional importance of 3D chromatin organization.

Chromatin interactions are potentially encoded in the DNA sequence through a sophisticated interplay of protein binding sites and various sequence elements. Many algorithms have been developed to predict genome folding based on its own sequence. DNA motifs associated with certain architectural proteins (such as CTCF proteins), promoters, enhancers, repetitive sequences, and noncoding RNA genes have been shown to play critical roles in 3D genome architecture (37–39). Considering that CTCF proteins are not described in trypanosomes and that the *T. cruzi* genome is composed of 50% repeats, where virulence factor genes constitute the majority, we aimed to evaluate the impact of different genomic features on chromatin folding and nuclear organization in *T. cruzi* by analyzing a Hi-C dataset.

## RESULTS

To assess the 3D genome structure of *T. cruzi*, we evaluated a Hi-C dataset characterizing TAD-like structures and their boundaries. First, we addressed missing genomic annotations of the *T. cruzi* BrazilA4 strain to investigate the genomic distribution of specific features (e.g., small rRNA genes, transcription initiation and termination sites, PTUs) across its 3D nuclear architecture.

### Improvements in the genome annotation, characterization of genomic compartments, and repetitive DNA per chromosome of the *T. cruzi* Brazil A4 strain

The available genome of the *T. cruzi* Brazil A4 strain (30) lacked annotations for snoRNAs, snRNAs, tRNAs, rRNAs, and spliced leader (SL) RNAs. Using an in-house pipeline, we identified 1199 snoRNA genes, 105 ncRNA (spliced-leader, SL, and uridine-rich small nuclear RNA (U-snRNA)) genes, and 201 rRNA (18S, 5S, 5.8S, and 24S) genes (Table S1). Additionally, we identified a total of 71 tRNA genes whose results proved to be as sensitive as the tRNAScan output, detecting the same number of tRNA genes at identical genomic positions. Moreover, our approach exhibits high precision, as we were able to identify four undetermined tRNA genes by tRNAScan, revealing them as valine tRNAs (Table S2). By identifying the orientation of both Box A and Box B for each tRNA gene, we generated a map of its distribution in the *T. cruzi* Brazil A4 genome, in which it is evident that tRNA genes tend to form clusters composed of two or more genes in arrays (Fig. S1). Additionally, we identified 1364 PTUs, accompanied by 407 cSSRs, 416 dSSRs, and 445 intergenic regions (IRs).

We computed the proportion, in terms of length, per chromosome of unidimensional genome organization (*core/disruptive/GpDR*) genes, repetitive DNA, and pseudogenes (Supplementary Figure 4). *Core* genes are more prevalent in the *T. cruzi Brazil A4* genome, which comprises 76.2% (11515 out of 15111 total genes), while *disruptive* and *GpDR* genes constitute 16.63% (2512 genes) and 7.17% (1084 genes) of the genome, respectively (Fig. S2B). Repetitive DNA is prevalent over the 43 chromosomes, particularly in those with a higher concentration of *GpDR* and *disruptive* genes (Fig. S2A and B). The top five repetitive-rich chromosomes are Chr38, 39, 35, 36, and 42 (Fig. S2C). Chromosomes with high repetitive DNA content usually have an increased number of pseudogenes (Fig. S2C). These comprehensive genomic annotations and characterizations played a pivotal role in our analysis of the 3D genome structure of *T. cruzi*, shown below.

### Topological associated domains (TAD)-like and chromatin loops reflect the 1D genomic organization of *core* and *disruptive* compartments

To evaluate the 3D genome structure of *T. cruzi*, we identified TAD-like domains using Hi-C matrices of various resolutions. We found that lower-resolution matrices resulted in fewer TAD-like regions with longer median lengths (Figure 1A; Fig. S3), suggesting a hierarchical organization with nested TAD-like domains (Fig. S3B). The median TAD length of 43 Kbp in 5 Kbp resolution Hi-C matrices (Fig. S3A) is significantly shorter than the 880 Kbp median TAD length observed in mammals (40), leading us to adopt the term “TAD-like domains” for *T. cruzi*. Higher resolution matrices (2-5 Kbp) lead to TAD-like domains that cover a greater percentage of the genome (68-79%) (Figure 1B), comparable to the 90% coverage in mammals (40). Therefore, we focused on matrices with 2-5 Kbp resolution for further analysis.

**Fig 1:**
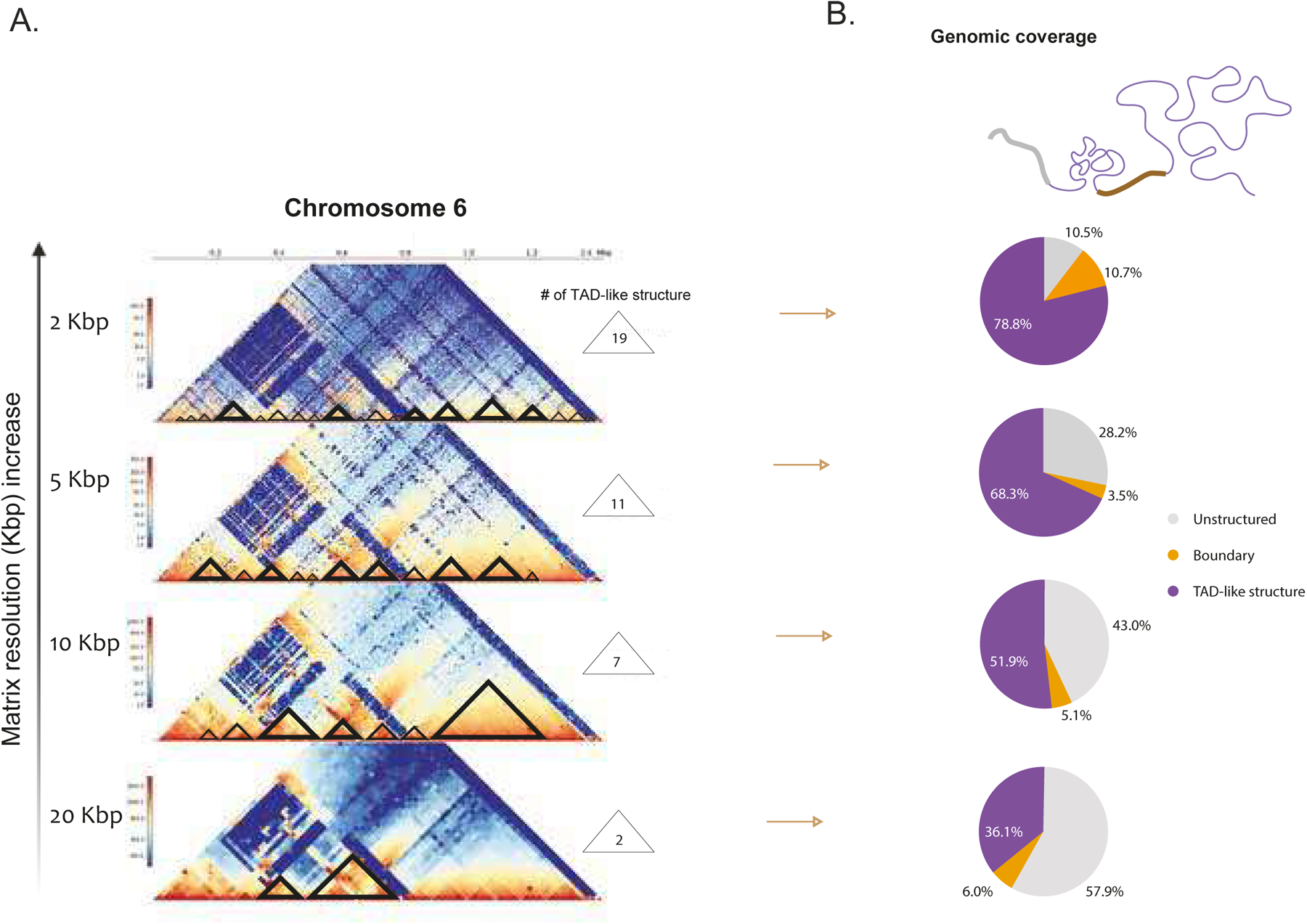
Hi-C matrix resolution effect in TAD-like calling of the *T. cruzi* Brazil A4 genome. A. Identification of TAD-like domains (black triangles - x-axis) in KR-normalized Hi-C heatmaps for chromosome 6 according to a matrix resolution ranging from 2 to 20 Kbp (y-axis). B. Pie charts representing the overall (43 chromosomes) genomic coverage percentages of the TAD-like domains, boundaries, and unstructured regions according to the matrix resolution.

We characterized TAD-like domains based on the unidimensional genomic composition of *T. cruzi* (*core* and *disruptive* compartments, in addition to *GpRD* genes). We examined whether these linear compartments also formed distinct 3D compartments. Our analysis revealed that 51% of the TAD-like domains were mixed (containing genes from more than one compartment), while 49% were pure (containing genes from only one compartment) (data not shown). Pure TAD-like domains composed of *core* genes were longer (4 to 178 Kbp) than those composed of *disruptive* and *GpDR* genes (4 to 28 Kbp) (Fig. S4). This variation in TAD-like length suggests that TADs-like enriched in *disruptive*/*GpDR* genes, which are shorter than the *core* TAD-like domains, are also more compact in the nucleus, as recently suggested (31).

To investigate the impact of these linear compartments on intrachromosomal interactions, we compared the number of chromatin loops in chromosomes enriched in the *core*, *disruptive* and *GpDR* genes. Notably, chromatin loops were more frequent in chromosomes enriched in *disruptive*/*GpDR* genes than in those enriched in core genes (Figure 2). For instance, chromosome 15, which is enriched in *core* genes (∼62.3%), had no loops, whereas chromosome 12, which is impoverished in *core* genes (∼17.3%), contained nine loops (Fig. 2A). This pattern was also observed for chromosome pairs 6 and 7, 12 and 13, and 25 and 26, which have similar lengths but differ in *core* gene enrichment (Fig. S4B). Chromosomes enriched in *core* genes (chrs 7, 13 and 26) had at least half the number of loops compared to those enriched in *disruptive* genes (chrs 6, 12 and 25) (Fig. 2B). Overall, we observed a negative correlation (R = -0.5) between the percentage of *core* genes and the number of loops across all 43 chromosomes (Fig. 2C). In summary, the TAD-like domains and chromatin loops in *T. cruzi* reflect the 1D genomic organization into *core* and *disruptive* compartments, influencing the 3D genome structure and interactions.

**Fig 2:**
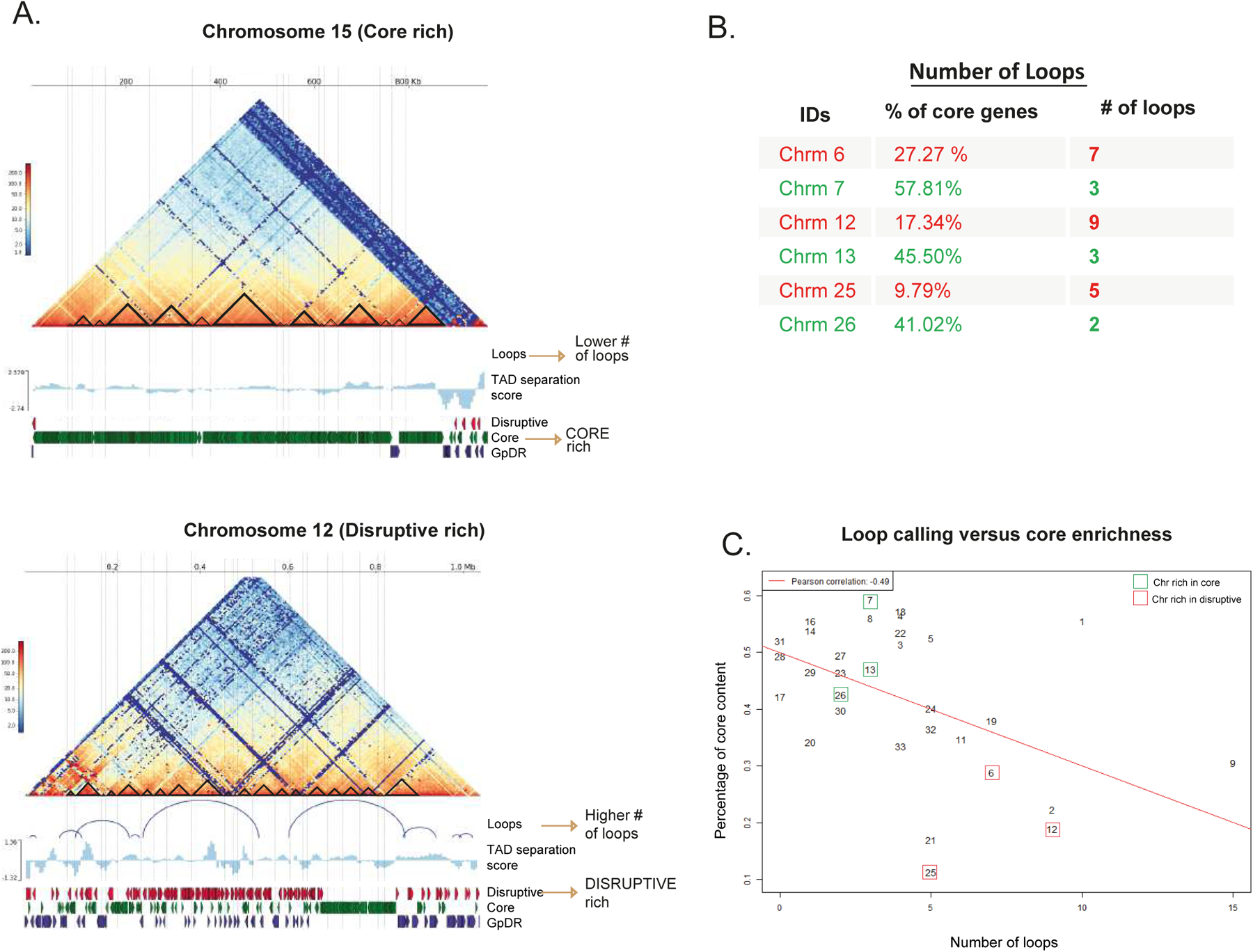
TAD-like domains enriched in *disruptive*/*GpDR* genes are shorter than TAD-like structures enriched in *core* genes, and *core*-rich chromosomes contain fewer DNA loops than *disruptive*/*<u>GpDR</u>*-rich ones. A. KR-normalized Hi-C heatmaps with the identification of loops (blue semicircles - x-axis) for a “*core*”-rich chromosome (chr 15) and a *disruptive* rich chromosome (chr 12). B. Analysis of chromosome pairs with differing *core* gene contents (e.g., chromosomes 6 and 7, 12 and 13 and 25 and 26). Chromosomes enriched in *core* genes (chrs 7, 13 and 26) displayed fewer loops than did those enriched in *disruptive* genes (chrs 6, 12 and 25). C. Dot plot of loop numbers (x-axis) per numbered chromosome. There was a negative correlation (R = -0.5) between the percentage of *core* genes and the number of chromatin loops across all 43 chromosomes, indicating that TAD-like domains and chromatin loops in *T. cruzi* are influenced by the linear genomic organization into *core* and *disruptive* compartments.

### Impact of multimapped reads detection on TAD-like identification

The *T. cruzi* genome, comprising more than 50% repetitive DNA (6), presents significant challenges for traditional Hi-C processing pipelines (41–43), which typically disregard multimapped reads. These reads are associated with low-quality mappings and multiple alignments, primarily occurring in repetitive genomic regions. To assess the impact of repetitive regions on 3D chromatin folding, we employed the multimapping strategy for Hi-C data (mHiC) pipeline (44), which optimizes the allocation of multimapped reads. We also compared the results with matrices obtained using the HiC Explorer tool, which excludes multimapped reads, a pipeline used in previous studies (31).

The inclusion of high-quality multimapped reads allowed us to allocate more than 12 million additional reads compared to traditional Hi‒C pipelines (Table S3). Discarding multimapped reads in traditional pipelines introduces gaps in Hi-C matrices (Figure 3A), which can obscure critical DNA‒DNA contacts and lead to erroneous conclusions. This issue is particularly pronounced in chromosomes enriched in *disruptive* genes, such as chromosomes 12 and 25, which consist of 72% and 86% repetitive DNA, respectively (Figure 3A). Fig. 3A illustrates the comparison between these two strategies, highlighting the improvements in gap filling, contact enhancement, and loop detection in four *T. cruzi* chromosomes (by mHi-C). This finding underscores the importance of including multimapped reads to reveal previously unknown DNA‒DNA contacts within the *T. cruzi* genome.

**Fig 3:**
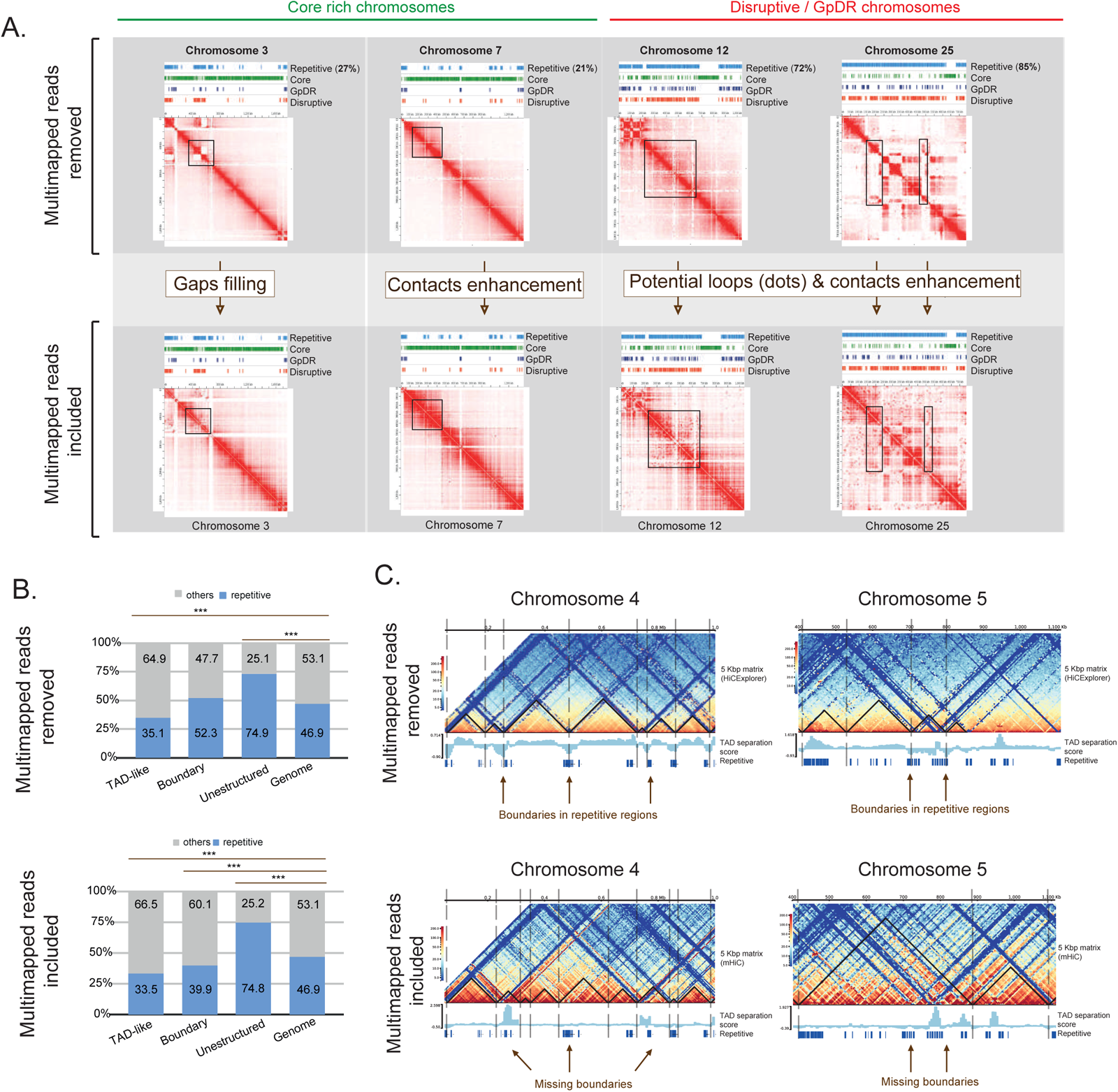
Repetitive DNA in Hi-C data analysis provides an enhancement of Hi-C contacts and avoids bias in calling TAD-like structures and their boundaries. A. Juicebox view of individual Hi‒C matrices obtained excluding multi-mapped reads via the Hi‒CExplorer pipeline (upper panel). The inclusion of multimapped reads from the mHi-C pipeline is shown in the lower panel. The black rectangles highlight DNA‒DNA contacts and loop enhancement, and gap filling recovered to the repetitive regions (light blue tracks). DNA loops are commonly seen in Hi-C matrices as punctual dots, potentially enriched for *disruptive* regions of the genome (Chr 15 and Chr 25). Tracks seen above the individual matrices stress multigenic families located in the *GpDR* (dark blue), *disruptive* (red), and *core* (green) compartments. B. Distribution of repetitive DNA (light blue) and nonrepetitive DNA (gray) within TAD-like domains, boundaries, unstructured regions and the genome (control) compared with the results obtained when processing Hi-C data excluding multimapped reads, HiCExplorer, upper panel, or including it, mHiC, lower panel, pipelines. *** p value < 0.01 using the chi-square test. C. Shifts in TAD boundaries (vertical dashed lines) and TAD-like domains (black triangles). The exclusion of repetitive reads showed that TAD-like boundaries were enriched in repetitive regions (e.g., chromosomes 4 and 5), while their inclusion shifted some boundaries to nonrepetitive regions, indicating biases in TAD calling due to the exclusion of multimapped reads.

Next, we evaluated the impact of repetitive regions on the identification of 3D genome structures, specifically TAD-like domains, their boundaries and unstructured regions. To achieve this goal, we compared the distribution of repetitive DNA within these structures (Figure 3B) against their expected distribution in the genome, which served as a control. Considering whether or not the reads are repetitive does not affect the identification of regions classified as unstructured, which harbor more than 70% repetitive DNA—a topic that will be explored later.

Excluding multimapped reads from the Hi-C data analysis resulted in the identification of 212 TAD-like domains and 250 boundaries (Fig. S6A). The inclusion of multiple mapped reads yielded similar numbers, with 203 TAD-like domains and 242 boundaries (Fig. S6A), as well as comparable median TAD length values (Fig. S6B). Notably, TAD-like domain boundaries exhibited approximately 52% enrichment in repetitive DNA when multimapped reads were excluded. Conversely, when multimapped reads were included, the repetitive DNA content at TAD-like boundaries decreased to approximately 40% (Figure 3B). Further examination of TAD-like boundaries across 20 chromosomes (Chr 1 to 20) confirmed that the inclusion of multimapped reads reduced the number of boundaries within repetitive regions and increased the percentage of boundaries detected in nonrepetitive DNA regions (Fig. S6C). Specifically, excluding multimapped reads highlighted TAD boundaries enriched in repetitive DNA on chromosomes 4 and 5. In contrast, including multimapped reads caused a shift, with certain boundaries previously identified in repetitive regions no longer evident (Fig. 3C). This result suggests that TAD calling is biased by the absence of Hi-C contacts in repetitive regions due to the exclusion of multimapped reads (e.g., using the HiCExplorer pipeline, 34% of the mapped reads were discarded; Table S3). Ultimately, these findings reinforce the importance of considering repetitive regions to accurately determine 3D genomic structure avoiding errouneous conclusions. Consequently, the analyses described below were performed with the included multimapped reads.

### TAD-like domains, boundaries and unstructured regions are characterized by distinct genomic sequences in *T. cruzi*

The comprehensive analysis of the 3D genome organization in *T. cruzi*, focusing on TAD-like domains and their boundaries, also involved evaluating the enrichment of specific genomic features within these structures. By comparing the observed versus expected distributions of certain loci in TAD-like domains, boundaries, and unstructured regions, we revealed several notable patterns. First, unstructured regions were enriched with genes from the MF compartment (*disruptive* and *GpDR* genomic compartments) but had fewer *core* genes (Figure 4A). MF genes are enriched in unstructured regions, coinciding with chromosome ends, as exemplified in chromosomes 5, 7 and 19 (Fig. S5). These regions are also enriched in pseudogenes (Figure 4B), aligning with the higher percentages of pseudogenes in the *GpDR* (82.7%) and *disruptive* compartments (61.5%) (Fig. S2B).

**Fig 4:**
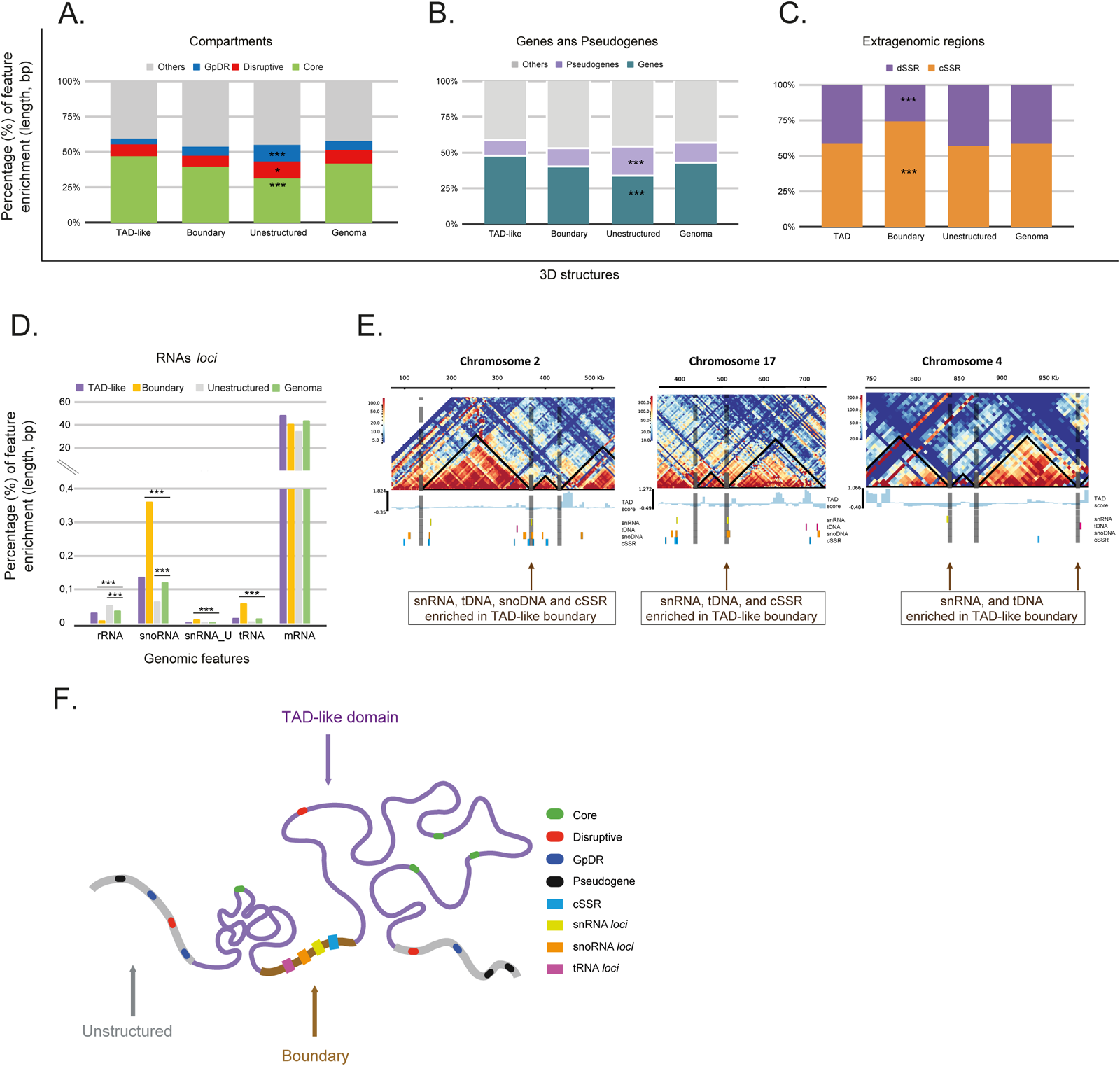
The nonrandom positioning of *T. cruzi* genomic features along TAD-like domains, their boundaries, and unstructured regions. A. Distribution profile of *core*, *disruptive*, and *GpDR* genes across 3D structures (TAD-like sctructures, boundaries and unstructured regions) compared to the genome as a control. *GpDR* (blue) genes and *disruptive* (red) compartments are enriched in unstructured regions. The others (gray) represent all of the genome sequences that remained after excluding the *core*, *disruptive* and *GpDR* sequences. B. Distribution of genes and pseudogenes. Pseudogenes (light purple) show a predominance in unstructured regions. The others (gray) represent all of the genome sequences that remained after excluding genes and pseudogenes. C. Distribution of extragenomic regions, specifically cSSR and dSSR sites, with cSSR sites predominantly located at boundaries. D. Distribution of small RNA genes (rRNA, snoRNA, snRNA_U, tRNA, and mRNA) across 3D genomic structures. Notably, snoRNA, snRNA_U, and tRNA loci are preferentially located at TAD-like boundaries. For A to D, the chi-square test indicates significance for p-adjusted values < 0.05. From A to D, the percentages represent the abundance of each feature (in length, bp) divided by the length of each 3D compartment or by the entire genome length for the controls. E. Enrichment of snRNAs, tRNAs, snoRNA genes, and cSSR at TAD-like boundaries for chromosomes 2, 17 and 4. F. Model illustrating the preferential positioning of target genomic features within the 3D nuclear architecture.

We asked whether each polycistronic units (PTUs) would be folded into a single or multiple TAD-like structures and vice versa. We identified three scenarios: PTUs folded into more than 1 TAD-like structure; PTUs formed by a unique TAD-like structure; and TAD-like structures covering more than 1 PTU. Considering that PTUs are delimitated by dSSRs and cSSRs, which demarcate the start and stop points of RNA polymerase II transcription, respectively, we evaluated their enrichment in 3D structures. In absolute numbers, 30 cSSRs (8%) and 17 dSSRs (5.5%) coincided with TAD-like boundaries. Thus, the majority of PTU borders (SSRs) do not coincide with TAD-like boundaries. However, considering that TAD-like boundaries cover only 3.5% (in length) of the genome, the presence of SSRs in these regions may be functionally relevant. Considering the relative length of each 3D structure to the entire genome, cSSRs were notably enriched at TAD boundaries and were 1.27 times more frequent than they were in the genome-wide distribution (Figure 4C). This result suggests that some boundaries may impact transcription initiation and termination.

We also investigated whether loci encoding various RNAs (mRNAs, tRNAs, snoRNAs, snRNAs, SL-RNAs, and rRNAs) were associated with specific 3D genome structures. We found significant enrichment of tRNA, snoRNA, and snRNA loci at TAD boundaries (Figure 4D). Considering only the group of RNA loci, the tRNA loci made up 7.7% of the genome by length and comprised 13.4% of TAD-like boundaries, indicating a 1.74-fold enrichment. Similarly, the enrichment of the snRNA and snoRNA loci at TAD-like boundaries was 1.17 and 1.85 times greater, respectively, than that at random genomic distribution. These findings were consistent across Hi-C matrices at a 10 Kbp resolution, underscoring the significance of these genomic features in the 3D organization of the *T. cruzi* genome (data not shown).

### The tRNA loci interactions suggest the formation of a spatial cluster within *T. cruzi* nuclei

Previously, it was found that the chromatin of tRNA loci is developmentally regulated (16). Here, we discovered that TAD-like boundaries are enriched in tRNA loci. To provide further insights into the role of tRNA loci in 3D genome organization, we used a virtual 4C approach to screen DNA‒DNA interactions mediated by all tRNA loci (viewpoints, VPs) (Figure 5). Remarkably, the interactions between tRNA loci are stronger than those between randomly selected control loci (tRNAs-*vs.*-controls, in green). This suggests that some tRNA loci form a 3D cluster through *cis-* and/or *trans*-acting interactions (Figure 5 – A, tRNA-*vs.-*tRNA panel). tRNA genes also exhibit a greater frequency of contact with snoRNAs, which are transcribed by RNA pol II (Figure 5A, tRNA-*vs.*-snoRNA panel). In contrast, significant interactions were not detected between tRNA loci and SL-RNA loci (also transcribed by RNA polymerase II) or between tRNA loci and rRNA loci (with 24S, 5.8S, and 18S rDNAs transcribed by RNA polymerase I and 5S rDNA genes transcribed by RNA polymerase III). snoRNAs and tRNA genes share a widespread genomic distribution, being the most prevalent non-protein-coding RNA loci across the *T. cruzi* genome, found on more than 12 different chromosomes (S1 Table). Both are enriched in TAD-like boundaries, suggesting that the interaction between tRNAs and snoRNAs may facilitate the grouping of TAD-like boundaries, promoting 3D contacts.

**Fig 5:**
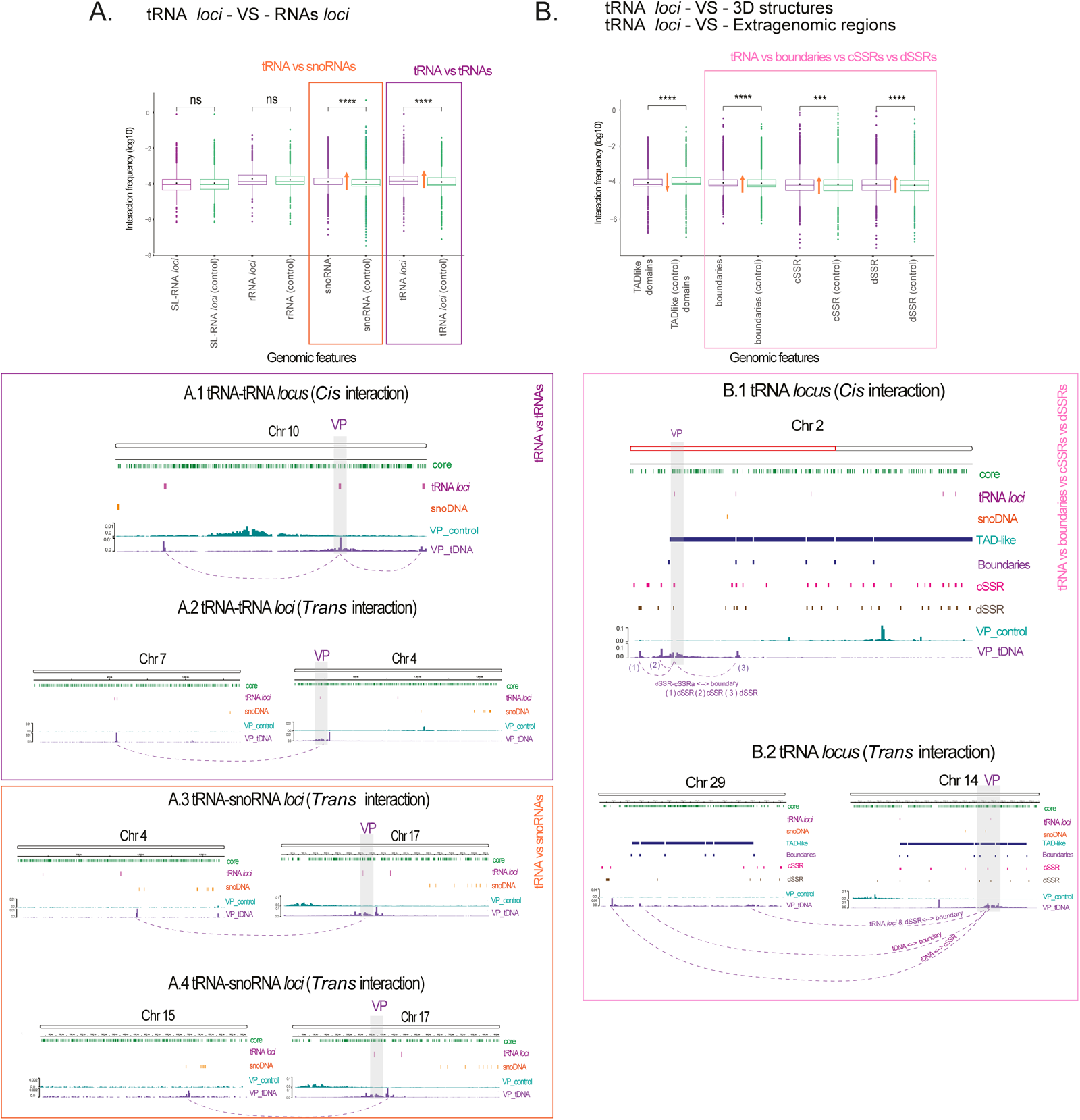
Virtual 4C analysis of tRNA loci as ViewPoints (VPs). Panels (A) and (B) display the comparison of DNA‒DNA interaction frequencies (log10) for tRNA loci with RNA loci and 3D genomic structure, respectively. For each target (purple boxplots), their control counterparts (green boxplots) are compared. Significant differences are indicated by *** p < 0.001 and **** p < 0.0001. Rectangles in orange, purple and rose indicate genomic features that have a greater frequency of contacts with tRNA genes than with their respective controls. The orange arrows indicate whether the interactions observed for VP are above or below the control interactions. A. Significant interactions between tRNAs and snoRNAs and between tRNAs and tRNAs are shown by orange and purple rectangles, respectively. Integrative Genomics Viewer (IGV) snapshots depicting *cis*-acting, intra, (A.1) and *trans*-acting, inter, (A.2) chromosomal contacts between the tRNA locus (VP) and other tRNA loci. *Trans*-acting (A.3 and A.4) interactions between the tRNA locus (VP) and snoRNA locus. B. Significant interactions highlight that DNA‒DNA contacts are frequent between tRNAs and TAD-like boundaries in addition to the extragenomic regions cSSRs and dSSRs (rose rectangle). *Cis*-acting (B.1) and *trans*-acting (B.2) chromosomal contacts among the tRNA locus (VP), 3D structures, and extra genomic regions.

In terms of 3D structures, tRNA loci preferentially interact with TAD-like boundaries and both dSSRs and cSSRs compared to the control regions used as VPs (Figure 5B). *Cis*-acting and *trans*-acting interactions represent DNA‒DNA contacts captured between tRNAs, 3D structures (boundaries) and extragenomic regions (cSSRs and dSSRs). Together, these results indicate that tRNA loci perform a vast set of 3D interactions, which include the following: the tRNAs themselves, the snoRNAs, TAD boundaries, cSSRs and dSSRs. Collectively, these results demonstrate that tRNA loci engage in a broad network of 3D interactions, highlighting their significance in the spatial organization of the *T. cruzi* genome.

### Remarked 3D nuclear networks involve the SL-RNA and 18S rRNA loci in *T. cruzi*

To better understand 3D local interactions within the nuclei, we extended the aforementioned analysis to other RNA loci, emphasizing SL-RNA and rRNA genes, which were both previously indicated as key genomic loci contributing to 3D nuclear organization (12, 45–47).

The SL-RNA locus, a highly transcribed gene essential for trans-splicing in *T. cruzi*, the unique gene transcribed by RNA polymerase II with promoter regions (2). Inspired by studies in *T. brucei* suggesting a role for SL-RNA loci in activating VSG genes through 3D proximity-based regulation (12), we assessed the 3D contact frequency for SL-RNA loci (VP on chromosome 23) with the rest of the genome by virtual 4C analysis (Figure 6A). The SL-RNA loci frequently interact with the 18S, 5.8S and 24S rRNA genes (found at chr 16) but not with the 5S rRNA loci (chr 8 and 29), as shown in the VP profiles of the rRNA genes relative to the SL-RNA loci at chr 23 (Fig. 6A).

**Fig 6:**
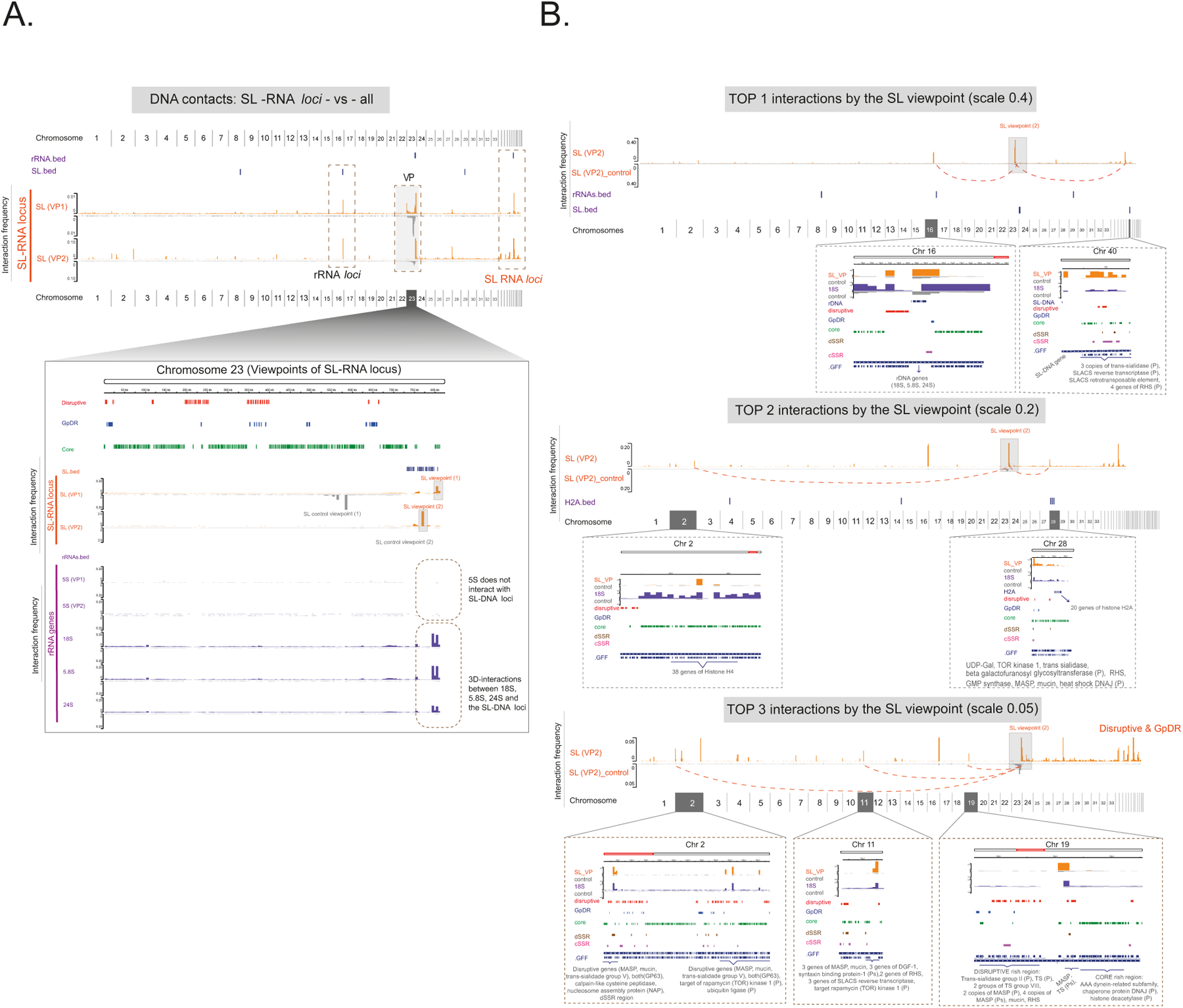
3D contacts involving the SL-RNA loci in *T. cruzi*. A. Virtual 4C plot profile. The plot shows interaction frequencies of SL-RNA loci (VPs), orange plot, across the entire genome (the x-axis represents the genomic coordinates to all 43 chromosomes). Control viewpoints are included for comparison via flipped gray plots. A zoomed-in view of chromosome 23 depicts the viewpoints (VP1 and VP2) of origin for the SL-RNA loci, highlighting the interaction frequencies with the 18S rRNA, 5.8S rRNA and 24S rRNA loci, excluding the 5S rRNA loci. B. DNA‒DNA interactions. The top 1 to top 3 plots focused on the greatest number of DNA‒DNA interactions between SL–RNA loci and other genomic regions, indicating significant 3D nuclear architecture features.

We identified seven main genomic regions exhibiting frequent interactions with the SL-RNA loci (Fig. 6B). The strongest interaction peaks included the rRNA locus (chr 16) and another SL-RNA locus (chr 40) (Fig. 6B). The second level (top 2) of frequent interactions comprised the histone H4 array (38 gene copies on chr 2), regions on chromosome 28 involving *disruptive/GpDR* genes (e.g., RHS, MASP, mucin), and conserved genes preceding the histone H2A array (20 gene copies) (Figure 6B). The third most frequent interactions (top 3), between the SL-RNA array and *disruptive/GpDR*-enriched genomic regions on chromosomes 2, 11 and 19 (Figure 6B, third ranking position), in addition to “*Disruptive* and *GpDR*” engagement, which are located on the latest chromosomes, represent SL interactions with genes from the compartment *disruptive* or the *GpDR* group, as shown in Fig. S7-peaks “d” to “h”.

The search for ribosomal RNA loci interactions was motivated by their critical role in the translation machinery and nucleolar organization (48). We investigated the 3D interactions of rRNA loci on chromosome 8 (68 gene copies) and chromosome 29 (1 gene copy) for the 5S rRNA loci and on chromosome 16 for the 18S, 5.8S and 24S rRNA gene arrays. The spatial network between the SL-RNA array (chr 23) and the 18S, 5.8S, and 24S loci, excluding the 5S RNA genes, is emphasized for chromosome 16 (Figure 7A).

**Fig 7:**
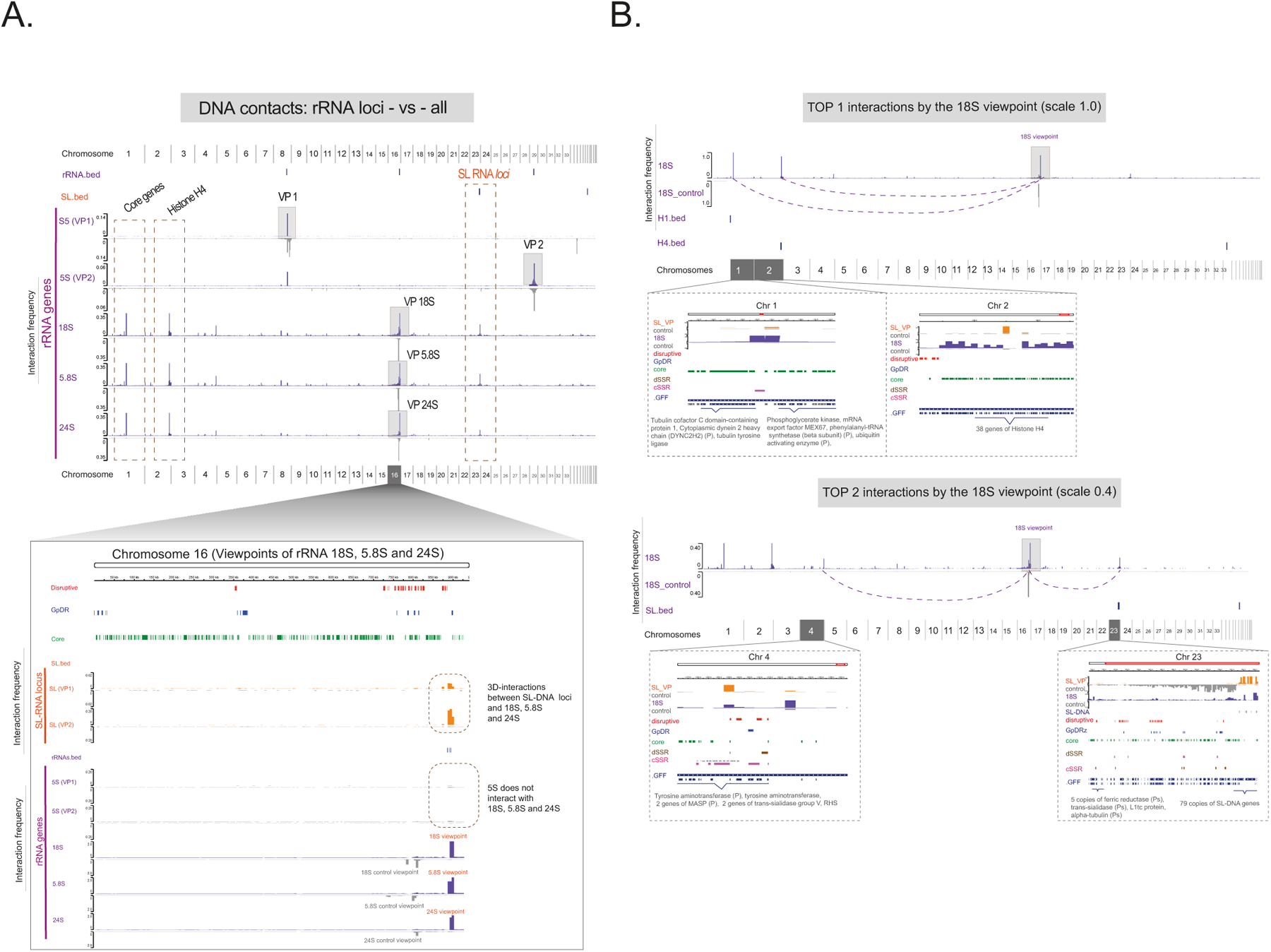
3D contacts involving the rRNA loci in *T. cruzi*. The plot profile highlights the virtual 4C results for the rRNA loci (VPs), purple plot, across the entire genome (the X-axis represents the genomic coordinates of all 43 chromosomes). Control viewpoints are included for comparison via flipped gray plots. A zoomed-in image of chromosome 16 shows the origin of the 18S rRNA, 5.8S rRNA and 24S rRNA loci. SL-RNA locus interaction sites are highlighted, and no remarkable interactions are observed between the three rRNA genes (18S, 5.8S, and 24S) or the 5S rRNA locus. B. The greatest number of DNA‒DNA interactions involving the 18S rRNA locus. The figure includes annotations for genomic elements such as *core* genes and histone H4 regions.

Given the similar whole-genome interaction profiles for all rRNA genes on chromosome 16, we selected the 18S viewpoint to inspect specific interactions (Figure 7B). The histone H4 locus (chr 2) showed a significant interaction frequency, sharing the top-ranking position with conserved genes on chromosome 1 involved in cell metabolism (e.g., phosphoglycerate kinase) and cytoskeleton assembly (e.g., tubulin- and dynein-related proteins). The second tier (top 2) of interactions includes the SL-RNA loci (chr 23) and an array of *disruptive/GpDR* genes (chr 4). Figure S10 compares the genome-wide interactions constrained by both the SL-RNA loci and 18S rRNA loci. The identified peaks are labeled A to Z (26 peaks) and a to h (8 peaks), totaling 34 peaks, with the exception of individual SL-RNA loci (3 peaks) or 18S rRNA loci (9 peaks) interactions. For those peaks exclusive to 18S rRNA loci, the 3D contacts to the *core* compartment are remarkable: cytoskeleton- and flagellum-related genes, heat shock genes (HSP70), and ribosomal subunits, among others (Fig. S7).

In turn, our 3D data indicate that *disruptive* and *GpDR* genes, which constitute approximately 17% and 7%, respectively, of all *T. cruzi* protein-coding genes (Fig. S2B – 1D view), interact more frequently with the SL-RNA loci (*Disruptive*/*GpDR*-*vs.*-SL-RNA) than with the 18S rRNA loci (*Disruptive/GpDR*-*vs.*-18S-rRNA) (Fig. 6 and Fig. S7 – 3D view). Overall, our findings illustrate a strong 3D linkage between two main loci, the rRNAs and the SL-RNA array (Figure 8). Additionally, we revealed a high frequency of contact of 18S rDNA loci (chromosome 16) and SL-RNA loci (chromosome 23) with the histone H4 array (chromosome 4) (Figures 6, 7). Mostly for the 18S rRNA locus VP, histones H1 (Fig. S7, peak C), H2B and H2BV (Fig. S7, peak I), and H2A (Fig. S7, peak b) were also engaged in *trans*-acting contacts. Together with the TAD-like structure data, loop calling, and genomic feature enrichment in 3D structures, we demonstrated the nonrandom 3D organization of *T. cruzi* nuclei (Figure 8).

**Fig 8:**
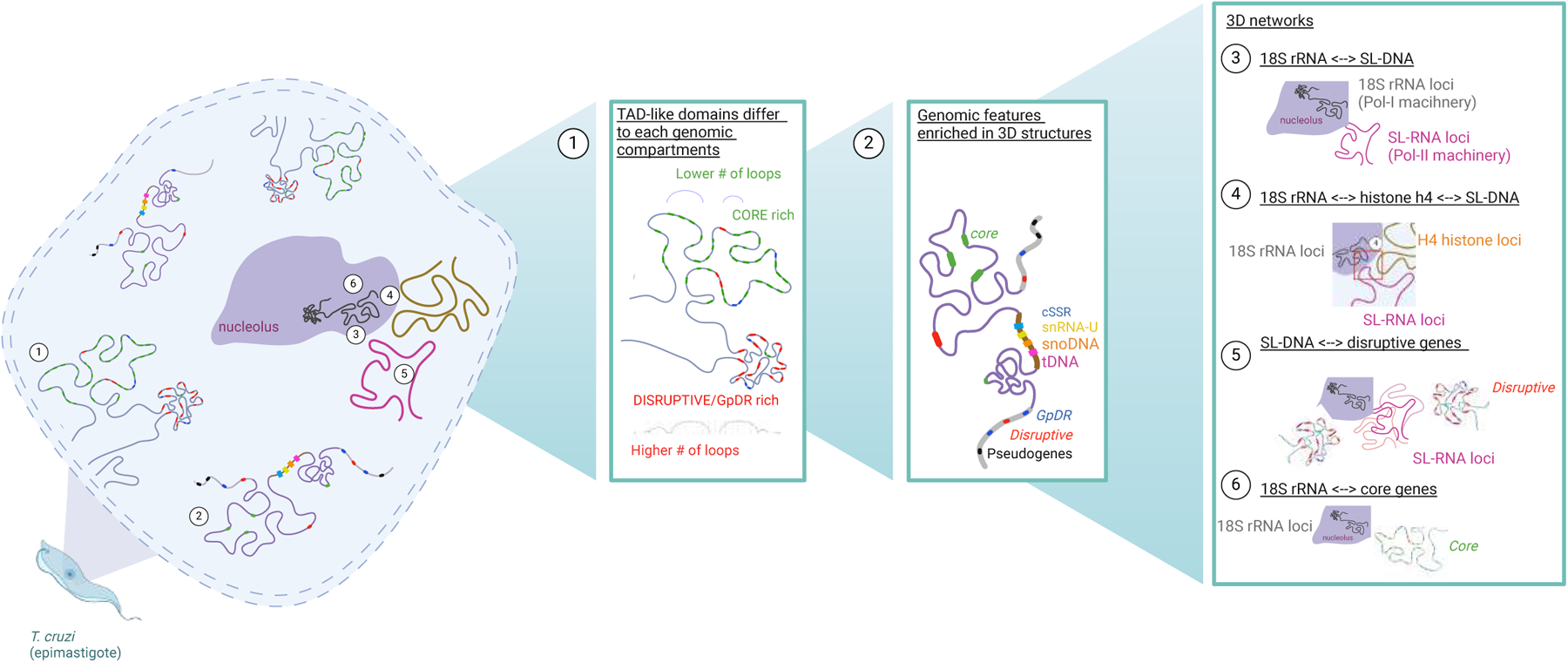
The nonrandom 3D nuclear architecture of *T. cruzi*. 1-Illustration of TAD-like domains and the formation of loops within the *core* (fewer loops) compartment and *disruptive*/*GpDR* genes (increased number of loops). 2-Genomic features relevant to 3D structure formation; 3-Nucleolar organization: Representation of the nucleolus, indicating the localization of 18S rRNA loci (Pol-I machinery) and SL-RNA loci (Pol-II machinery). 4-DNA‒DNA interactions for rRNA loci, the histone H4 array, and the SL–RNA loci; 5-Remarkable networks within *T. cruzi* nuclei reveal frequent interactions for SL–RNA loci and *disruptive*/*GpDR* genes, 6-while for the 18S rRNA loci, the *core* genes often interact. Model built with BioRender.

## DISCUSSION

The *T. cruzi* genome, characterized by its high repetitive content (51.2%) (49), presents challenges for Hi-C read mapping due to the prevalence of repetitive regions that generate low-quality mappings, which are often withdrawn during Hi-C processing. To overcome this issue, we used the mHi-C pipeline (44) to recover Hi-C contacts for genomic regions enriched in repetitive DNA. This strategy allowed comprehensive mapping and analysis of the 3D chromatin organization patterns of *T. cruzi* while also mitigating biases in the identification of TAD-like domains, their boundaries, and the participation of surface multigenic families (MFs).

The MF genes mucins, MASPs, and trans-sialidases play essential roles in cell invasion and immune response modulation during infection (50), and constitute approximately 29% of the *T. cruzi* repeatome (49). We observed that the MF genes, which exhibited lower transcriptional activity in nonreplicative forms, formed shorter and more compact TAD-like domains, while the *core* genes, with higher transcription rates and open chromatin levels in the epimastigote stage (51), formed longer TAD-like domains. Associations between the size of TAD-like structures and genomic compartments have been previously reported (31), but these 3D regions were better elucidated here with the inclusion of multimapped regions. The distinct genomic arrangement of these compartments is correlated with distinct gene activity levels and open chromatin profiles (16, 31, 51).

Aside from the TAD-like’s size, our data indicated that chromosomes enriched in *core* genes had fewer chromatin loops than did those enriched in *disruptive/GpDR* genes, which had at least twice as many loops, potentially associated with a gene regulatory mechanism. This organization mirrors *T. brucei* findings: a lower DNA‒DNA contact frequency in core gene regions and a greater frequency in subtelomeric regions (rich in VSG expression sites), indicating greater chromatin compaction in VSG-enriched areas (52). In *T. cruzi,* MF genes are dispersed throughout the genome (1D view) and preferentially occupy unstructured 3D genomic regions. We hypothesize that certain segments of DNA may adopt a 3D conformation to safeguard loci that may not be essential during a specific life stage but could play crucial roles in other physiological scenarios. Therefore, further research is needed to better understand the relevance of this spatial organization to parasite biology.

The three-dimensional folding of chromatin is determined by various factors encoded in the linear genome. These include repetitive DNA elements, as well as DNA motifs for architectural proteins, promoters and enhancers that form chromatin loops essential for gene regulation, and noncoding RNA (ncRNA) loci (38, 53–55). Together, these elements help organize chromatin by connecting different genome regions and shaping the intricate 3D structure of chromatin, impacting genomic architecture and function. In human and mouse genomes, tRNA loci mark TAD boundaries (40). In *Drosophila melanogaster*, these DNA motifs and the open chromatin state define 3D structures (56). In *T. brucei*, the cohesin subunit SSC1 is enriched in tRNA genes and cSSRs (29). tRNA loci were previously shown to be present at some TAD-like boundaries in *T. cruzi* (31). Here, we demonstrated that TAD-like boundaries are also proportionally enriched (in length) in noncoding RNA loci, such as tRNAs, snoRNAs, and snRNAs, along with cSSRs. In *T. cruzi*, tRNA and snRNA loci are linked to more open chromatin regions, especially in epimastigotes (replicative life form) than in trypomastigotes (infective form) (16). TAD-like boundaries are regions that restrict interactions of *cis*-regulatory sequences, contributing to the gene expression regulation of genes within TADs (40, 57). Although the PTU borders do not coincide perfectly with the TAD-like boundaries, those that coincide may be functionally relevant. For those case, it remains to be evaluated whether PTUs located in the same TAD have similar transcription rates, since we detected variations in PTU transcription, particularly between PTUs from different genomic compartments (51).

TAD boundaries are enriched in DNA motifs for insulator proteins such as Boundary Element Associated Factor-32 (BEAF-32) and CCCTC-binding factor (CTCF) in flies and mammals, respectively (56, 58, 59). CTCT is a core architectural protein that establishes the 3D structure of many eukaryotic genomes (60). Since trypanosomes lack CTCF proteins, the identification of TAD boundaries provided by our work has the potential to support further studies devoted to identifying regulatory proteins involved in 3D chromatin folding. This finding raises the possibility that trypanosomes harbor special alternative mechanisms to the well-known CTCF-based scheme encountered in most metazoans. Using the origami analogy, CTCF, cohesin and similar proteins act like human fingers, precisely folding chromatin fibers into specific shapes.

Our data revealed that tRNA loci preferentially interact with one another in a 3D context, suggesting the formation of clusters that likely optimize transcription by RNA polymerase III, similar to what is observed in yeast (61). The improved *T. cruzi* genome annotation led to the 1D mapping of tRNA genes, highlighting clusters with two to seven genes in tandem, corroborating our hypothesis that RNA polymerase III can be used to optimize transcription. We identified a recurrent match for the 3D positioning of tDNAs with cSSRs, dSSRs, snoRNAs and boundaries of TAD-like domains, which together suggest the regulation of the transcriptional machinery of RNA polymerase II and RNA polymerase III, as already claimed for yeast (62).

The SL-RNA loci interact 3D predominantly with rRNA loci, with the histone H4 gene cluster, and with trans-sialidases, mucins, RHS, genes and pseudogenes (P) from the *disruptive/GpDR*-enriched genomic sites. The interactions with MF genes coding for virulence factors suggest a potential regulatory role of the SL-RNA loci for these regions, resembling its role as a putative posttranscriptional enhancer of monoallelic VSG expression in *T. brucei* (12). Notably, VSGs are transcribed by RNA Pol I, while *T. cruzi* MFs are transcribed by RNA Pol II. Therefore, we hypothesize that virulence factor expression in *T. cruzi* may take advantage of the proximity of the RNA Pol II machinery to boost both posttranscriptional and transcriptional mechanisms. Further investigations evaluating whether these 3D structures change in *T. cruzi* infective forms must be performed to better understand their impact on gene expression.

The nucleolus is a key subnuclear structure involved in rRNA synthesis and ribosome assembly (63) and plays a key role in the organization of 3D genome architecture. In mammals, nucleolus-associated chromatin interactions form strong heterochromatin interactions (46), and nucleolar-associated domains (NADs) are AT-rich DNA sequences (47). Our study revealed that in *T. cruzi*, the 18S rRNA interactions include mainly *core* genes, which exhibit lower GC content (7). These *core* interactions include genes related to the cytoskeleton and flagellum, histone genes (H4, H1, H2B, H2BV, and H2A), ribosomal structural components, and heat shock genes such as HSP70 and snoRNA genes (66 copies). These interactions suggest that *T. cruzi* nucleoli interactions include transcribed/euchromatin regions (core genes) (51, 64), in contrast to other eukaryotes. Nevertheless, the *T. cruzi* nucleolus extends its role beyond ribosomal biogenesis by shaping the 3D chromatin structure, as observed in other eukaryotes (13). In *T. brucei*, the SL locus transcription compartment associated with active VSG is also closer to the nucleolus (12). In *T. cruzi*, RNA polymerase II is concentrated near nucleoli and enriched at SL-RNA genes, indicating clustering of pol I and pol II RNA polymerases (45). Our findings confirm this previous observation showing significant spatial contact between rRNA loci and SL-RNA loci. Interestingly, the 5S rRNA gene, transcribed by RNA Pol III, is distributed separately from the 18S rDNA/SL-DNA network, consistent with previous findings (11).

In addition to interacting with each other, we found that 18S rRNA genes prioritize interactions with *core* genes, while the SL-RNA loci, although they are associated with some *core* genes, interact mostly with *disruptive*/*GpDR* genes. We hypothesize that the disassembly of the nucleolus in infective forms (22) may affect these interactions, likely affecting their expression.

The role of chromatin architecture in transcription control has been well documented across various organisms (65, 66). Previously, we found that the open chromatin status and nucleosome deposition were associated with mature (15, 16) and nascent transcript levels (51). Recently, the boundaries of TADs were linked to a lack of mature RNA, leading to the assumption that trypanosome gene expression is determined by 3D organization (31). Here, we provide new insights into the role of repeated regions, which have been demonstrated to be crucial for the determination of TAD boundaries. Our results suggest that the 3D nuclear architecture of *T. cruzi* might impact gene expression regulation through spatial interactions among different genomic loci, including the coordination of gene expression involving genomic loci transcribed by distinct RNA polymerases. The existence of 3D chromatin interactions may optimize cross-talk among RNA polymerase machineries, transcription factors and epigenetic marks, which maintain or facilitate the transcription of target genomic regions.

## MATERIALS AND METHODS

### Data collection and read filtering

A Hi-C public dataset from *T. cruzi,* Brazil A4 strain, available at the National Center for Biotechnology Information (NCBI) repository (SRA number SRX8355434) (30) was used in this work to characterize the 3D nuclear architecture profile. The quality of the paired-end raw Illumina FASTQ reads was first assessed using FastQC v0.12.1 (https://www.bioinformatics.babraham.ac.uk/projects/fastqc). The trimming of the adaptors from the forward and reverse sequenced reads, removal of low-quality nucleotides at 5’, and removal of reads shorter than 70 bp in length were performed using Trimmomatic v.0.39 (67). The downstream data analysis was performed using the Hi-C Explorer pipeline v.3.0. (41), which entails a comprehensive suite of scripts for mapping, preprocessing (normalization), analysis, and visualization of the Hi-C data.

### Mapping

Filtered R1 and R2 FASTQ files were mapped against the *T. cruzi* Brazil A4 reference genome (GenBank access GCA_015033625.1, version NCBI 2020), which was previously digested in silico using the hicFindRestSite tool with “GATC” as the cutoff site to identify DpnII restriction sites. We employed the Burrows‒Wheeler Aligner (BWA) for mapping following the 4D Nucleome guidelines (https://data.4dnucleome.org/resources/data-analysis/hi_c-processing-pipeline).

### Hi-C data preprocessing

The R1 and R2.sam files generated during mapping were utilized for contact matrix construction via the hicBuildMatrix tool at a 0.5 Kbp bin resolution, followed by the merging of neighboring matrix bins to obtain lower resolutions ranging from 2 to 20 Kbp (the hicMergeMatrixBins tool). Normalization of all Hi-C pairwise matrices was performed using the Knight-Ruiz (KR) method (68) to mitigate biases such as DNA compaction range, variations in GC content, and fluctuations in copy number.

### General Feature Format (GFF) improvement

#### Nonprotein coding RNA genes (ncRNA genes)

Identification of small RNA genes was performed by BLASTn searches using the *T. cruzi* Dm28c genome-release54 (http://tritrypdb.org/) as the query, facilitated by a custom bash script called “findbestmatch” (available at: https://github.com/trypchromics/tcruzi-origami). This script constructed BLAST indices for the *T. cruzi* Brazil A4 genome, executed BLASTn searches, and processed the .csv results to identify the best matches. Our analysis of the transferred RNA (tRNA) genes was compared with the tRNAScan web server (69). Additionally, a manual BLASTn search was conducted specifically for the selenocysteine tRNA gene using homologous sequences from *T. brucei* (Tb927.9.2380 and Tb27.9.2340). Box A and Box B sites were then identified for all tRNA genes using the strategy proposed by (70). The “findbestmatch” script was also utilized for the identification of small nucleolar RNAs (snoRNAs), small nuclear RNAs (snRNAs), spliced-leader RNAs (SL-RNAs), and ribosomal RNAs (rRNAs). *<u>Extragenomic regions:</u>* Annotation of polycistronic transcription units (PTUs), transcription initiation sites (TSSs or dSSRs), termination sites (TTSs or cSSRs), and intergenic regions (IRs) was performed with the python-based “annotatePolycistron” script (https://github.com/alexranieri/annotatePolycistron).

### Assignment of target genes

#### Core/disruptive/GpDR

The GFF of *T. cruzi* Brazil A4 (NCBI version 2020) was screened to identify *core* and *disruptive* genomic compartments. Mucins, mucin-associated proteins (MASP), and trans-sialidase genes (TS) were filtered out and designated as the *disruptive* .GFF file. The GP63, dispersed gene family 1 (DGF-1), and retrotransposon hot spot (RHS) genes composed the GpDR .GFF file. The remaining filtered genes composed the *core* (housekeeping genes) .GFF. These files were further subdivided into “genes” and “pseudogenes”. All of these groups were converted into .BED files for visualization with Integrative Genomics Viewer (IGV), and the percentage of target genes per chromosome was determined using the custom script “bedCoveragePerChrom” (available at https://github.com/trypchromics/tcruzi-origami). *<u>Repetitive DNA:</u>* Based on the repeat list provided by (30), repeats shorter than 1000 bp were excluded, resulting in 5390 repetitive sequences grouped as DNAs (e.g., DNA/CMC-EnSpm, DNA/Zisupton), LINEs (e.g., LINE/CR1-Zenon, LINE/Jockey), low complexity A-rich regions, LTRs, rRNAs, simple repeats, satellites, and unknown types.

### TAD-like calling

TAD-like domains and boundaries were identified using the hicFindTAD tool from HiCexplorer v.3.0. To determine the optimal settings for the *T. cruzi* genome (approximately 40 Mbp), we tested our 2Kbp and 5Kbp matrices with deltas and thresholds ranging from 0.005 to 0.5. After evaluation, we selected a delta and threshold of 0.01 for both, as this configuration minimized spurious TAD identification (data not shown).

### Chromatin loop calling

Chromatin loops were identified using the hicDetectLoops tool (71). Loop calling was conducted at a 2 Kbp resolution to compare loops in *disruptive*-rich chromosomes versus *core*-rich chromosomes.

### mHi-C tool

The mHiC scripts (https://github.com/yezhengSTAT/mHiC) were adapted to suit the *T. cruzi* context. This tool involves two mapping steps, both of which are performed using the BWA aligner with parameters set according to the 4DN guidelines. After the first mapping round, high-quality multimapping reads from the .sam files were recovered, trimmed at the left 5’, and remapped onto the reference genome. Invalid Hi-C reads (e.g., self-circles, short-range interactions, self-ligations, dangling ends, and others of unknown origin, referred to as dump reads) were removed. Data normalization was performed using the Knight-Ruiz (KR) algorithm. The multimapping reads were then converted into uniquely mapped reads, and the final mHi-C outputs were generated as binary .mHiC files. The “.mHiC” matrices were converted into “.hic” files by an in-house script to enable their visualization with the JuiceBox interactive Hi-C contact matrix viewer (https://aidenlab.org/juicebox/).

### 3D higher-order interactions between tRNA loci, rRNAs, and SL-RNA loci (Virtual4C)

The 4C approach, which uses the 3C principle combined with high-throughput sequencing, investigates DNA‒DNA interactions from a unique locus of interest (the viewpoint, VP) against the entire genome (72). Its virtual version, named virtual 4C, derives one-versus-all interactions from the genome-wide Hi-C matrix. In this study, the genomic coordinates of all tRNA genes formed 31 VPs after excluding overlaps (regions where a set of tRNA genes are arranged with less than 2 Kbp between each other). These regions were individually examined as VPs and input into the HiC Sunt Dracones (HiCSD) package (https://github.com/foerstner-lab/HiCsuntdracones). The pipeline outputs a .wig file recording the set of Hi-C interactions constrained by each VP. The controls consisted of 31 genomic segments of similar size (bp) to the tRNA genes but located in different genomic regions. These genes were randomly sampled from the genome using the bedtools package, random function, with the options -l 73 (length), -n “variable” (number of control regions per chromosome, based on the number of real tRNA VPs per chromosome), and -seed 7135 (for shuffling). To determine the most likely 3D interactions, we calculated the mean Hi-C interactions for all tRNA genes, focusing on target genomic features (e.g., RNA loci, TAD-like domains, boundaries, and extragenomic regions). An R script, based on the bedtools coverage function, was customized to calculate tRNA interactions with each target region. The virtual 4C signals were plotted and compared to those of random controls. The same method was applied to rRNA genes and SL-RNA loci for which we directly evaluated the .wig files, which were plotted in comparison to their corresponding control VPs.

## ACKNOWLEDGMENTS

We thank Dr. Nicolai Siegel, Ms. Claudia Rabuffo, Dr. Markus Schmidt and Dr. Ye Zheng for supporting the settling of the mHiC code for the *T. cruzi* genome and very helpful suggestions. We thank Mariana Loterio Silva and Letícia de Sousa Lopes for carefully reading of the manuscript, as well as all TrypChrOMICś lab members for the many insightful discussions.

## REFERENCES

1. Lidani KCF, Andrade FA, Bavia L, Damasceno FS, Beltrame MH, Messias-Reason IJ, Sandri TL. 2019. Chagas disease: From discovery to a worldwide health problem. J Phys Oceanogr 49:1–13.

2. Gilinger G, Bellofatto V. 2001. Trypanosome spliced leader RNA genes contain the first identified RNA polymerase II gene promoter in these organisms. Nucleic Acids Res 29:1556–1564.

3. Teixeira SM, de Paiva RMC, Kangussu-Marcolino MM, DaRocha WD. 2012. Trypanosomatid comparative genomics: Contributions to the study of parasite biology and different parasitic diseases. Genet Mol Biol 35:1–17.

4. Clayton C. 2019. Regulation of gene expression in trypanosomatids: Living with polycistronic transcription. Open Biol 9.

5. Herreros-Cabello A, Callejas-Hernández F, Gironès N, Fresno M. 2020. *Trypanosoma cruzi* Genome: Organization, Multi-Gene Families, Transcription, and Biological Implications. Genes (Basel) 11.

6. El-sayed NM, Myler PJ, Bartholomeu DC, Nilsson D, Aggarwal G, Westenberger SJ, Tran A, Ghedin E, Worthey EA, Delcher AL, Caler E, Cerqueira GC, Branche C, Haas B, Anupama A, Arner E, Lena A, Burton P, Cadag E, Campbell DA, Attipoe P, Bontempi E, Carrington M, Crabtree J, Darban H, Franco J, Jong P De, Frasch AC, Gull K, Horn D, Hou L, Huang Y, Kindlund E, Klingbeil M, Kluge S, Koo H, Lacerda D, Levin MJ, Lorenzi H, Louie T, Machado CR, Mcculloch R, Mckenna A, Mizuno Y, Mottram JC, Nelson S, Ochaya S, Osoegawa K, Pai G, Parsons M, Pentony M, Pettersson U, Pop M, Ramirez JL, Rinta J, Robertson L, Salzberg SL, Sanchez DO, Seyler A, Sharma R, Shetty J, Simpson AJ, Sisk E, Tammi MT, Tarleton R, Teixeira S, Aken S Van, Vogt C. 2005. The Genome Sequence of *Trypanosoma cruzi*, Etiologic Agent of Chagas Disease. Science (80-) 4975:409–416.

7. Berná L, Rodriguez M, Chiribao ML, Parodi-Talice A, Pita S, Rijo G, Alvarez-Valin F, Robello C. 2018. Expanding an expanded genome: long-read sequencing of *Trypanosoma cruzi*. Microb genomics 4:1–19.

8. Nascimento J de F, Kelly S, Sunter J, Carrington M. 2018. Codon choice directs constitutive mRNA levels in trypanosomes. Elife 7:1–26.

9. Jeacock L, Faria J, Horn D. 2018. Codon usage bias controls mRNA and protein abundance in trypanosomatids. Elife 7:1–20.

10. Michaeli S. 2011. Trans-splicing in trypanosomes: Machinery and its impact on the parasite transcriptome. Future Microbiol 6:459–474.

11. Martínez-Calvillo S, Florencio-Martínez LE, Nepomuceno-Mejía T. 2019. Nucleolar structure and function in trypanosomatid protozoa. Cells 8.

12. Faria J, Luzak V, Müller LSM, Brink BG, Hutchinson S, Glover L, Horn D, Siegel TN. 2021. Spatial integration of transcription and splicing in a dedicated compartment sustains monogenic antigen expression in African trypanosomes. Nat Microbiol 6:289–300.

13. Symonová R. 2019. Integrative rDNAomics-importance of the oldest repetitive fraction of the eukaryote genome. Genes (Basel) 10:1–15.

14. Pontvianne F, Carpentier M-C, Durut N, Pavlištová V, Jaške K, Schořová Š, Parrinello H, Rohmer M, Pikaard CS, Fojtová M, Fajkus J, Sáez-Vásquez J. 2016. Identification of Nucleolus-Associated Chromatin Domains Reveals a Role for the Nucleolus in 3D Organization of the A. thaliana Genome.Pontvianne, F., Carpentier, M.-C., Durut, N., Pavlištová, V., Jaške, K., Schořová, Š., Parrinello, H., Rohmer, M., Pikaa. Cell Rep 16:1574–1587.

15. Lima ARJ, de Araujo CB, Bispo S, Patané J, Silber AM, Elias MC, da Cunha JPC. 2021. Nucleosome landscape reflects phenotypic differences in *Trypanosoma cruzi* life forms. PLoS Pathog 17:1–26.

16. Lima ARJ, Silva HG de S, Poubel S, Rosón JN, de Lima LPO, Costa-Silva HM, Gonçalves CS, Galante PAF, Holetz F, Motta MCMM, Silber AM, Elias MC, da Cunha JPC. 2022. Open chromatin analysis in *Trypanosoma cruzi* life forms highlights critical differences in genomic compartments and developmental regulation at tDNA loci. Epigenetics Chromatin 15:1–16.

17. Reynolds DL, Hofmeister BT, Cliffe L, Siegel TN, Anderson BA, Beverley SM, Schmitz RJ, Sabatini R. 2016. Base J represses genes at the end of polycistronic gene clusters in Leishmania major by promoting RNAP II termination. Mol Microbiol 101:559–574.

18. Jensen BC, Phan IQ, McDonald JR, Sur A, Gillespie MA, Ranish JA, Parsons M, Myler PJ. 2021. Chromatin-Associated Protein Complexes Link DNA Base J and Transcription Termination in Leishmania. mSphere 6.

19. Rosón JN, De Oliveira Vitarelli M, Costa-Silva HM, Pereira KS, Da Silva Pires D, De Sousa Lopes L, Cordeiro B, Kraus AJ, CruzI KNT, Calderano SG, Fragoso SP, Nicolai Siegel T, Elias MC, Da Cunha JPC. 2022. H2B.V demarcates divergent strand-switch regions, some tDNA loci, and genome compartments in *Trypanosoma cruzi* and affects parasite differentiation and host cell invasion. PLoS Pathog 18:1–27.

20. Siegel TN, Hekstra DR, Kemp LE, Figueiredo LM, Lowell JE, Fenyo D, Wang X, Dewell S, Cross GAM. 2009. Four histone variants mark the boundaries of polycistronic transcription units in Trypanosoma brucei. Genes Dev 23:1063–1076.

21. Kraus AJ, Vanselow JT, Lamer S, Brink BG, Schlosser A, Siegel TN. 2020. Distinct roles for H4 and H2A.Z acetylation in RNA transcription in African trypanosomes. Nat Commun 11.

22. Elias MCQB, Marques-Porto R, Freymüller E, Schenkman S. 2001. Transcription rate modulation through the *Trypanosoma cruzi* life cycle occurs in parallel with changes in nuclear organisation. Mol Biochem Parasitol 112:79–90.

23. Marstrand TT, Storey JD. 2014. Identifying and mapping cell-type-specific chromatin programming of gene expression. Proc Natl Acad Sci U S A 111.

24. Lieberman-aiden E, Berkum NL Van, Williams L, Imakaev M, Ragoczy T, Telling A, Amit I, Lajoie BR, Sabo PJ, Dorschner MO, Sandstrom R, Bernstein B, Bender MA, Groudine M, Gnirke A, Stamatoyannopoulos J, Mirny LA. 2009. Comprehensive Mapping of Long-Range Interactions Reveals Folding Principles of the Human Genome. Science (80-) 33292:289–294.

25. Galganski L, Urbanek MO, Krzyzosiak WJ. 2017. Nuclear speckles: Molecular organization, biological function and role in disease. Nucleic Acids Res 45:10350–10368.

26. Mitrentsi I, Lou J, Kerjouan A, Verigos J, Reina-San-Martin B, Hinde E, Soutoglou E. 2022. Heterochromatic repeat clustering imposes a physical barrier on homologous recombination to prevent chromosomal translocations. Mol Cell 82:2132–2147.e6.

27. Dekker J, Rippe K, Dekker M, Kleckner N. 2002. Capturing chromosome conformation. Science (80-) 2157:1–7.

28. Cullen KE, Kladde MP, Seyfred MA. 1993. Interaction between transcription regulatory regions of prolactin chromatin. Science (80-) 261:203–206.

29. Müller LSM, Cosentino RO, Förstner KU, Guizetti J, Wedel C, Kaplan N, Janzen CJ, Arampatzi P, Vogel J, Steinbiss S, Otto TD, Saliba AE, Sebra RP, Siegel TN. 2018. Genome organization and DNA accessibility control antigenic variation in trypanosomes. Nature 563:121–125.

30. Wang W, Peng D, Baptista RP, Li Y, Kissinger JC, Tarleton RL. 2021. Strain-specific genome evolution in *Trypanosoma cruzi*, the agent of Chagas disease. PLoS Pathog 17:1–30.

31. Díaz-Viraqué F, Chiribao ML, Libisch MG, Robello C. 2023. Genome-wide chromatin interaction map for *Trypanosoma cruzi*. Nat Microbiol 8:2103–2114.

32. Gibcus JH, Dekker J. 2013. The Hierarchy of the 3D Genome. Mol Cell 49:773–782.

33. Ye C, Paccanaro A, Gerstein M, Yan KK. 2020. The corrected gene proximity map for analyzing the 3D genome organization using Hi-C data. BMC Bioinformatics 21:1–18.

34. Beagrie RA, Scialdone A, Schueler M, Dorothee CA. 2017. Complex multi-enhancer contacts captured by Genome Architecture Mapping (GAM). Nature 543:519–524.

35. Fullwood MJ, Liu MH, Pan YF, Liu J, Xu H, Mohamed Y Bin, Orlov YL, Velkov S, Ho A, Mei PH, Chew EGY, Huang PYH, Welboren WJ, Han Y, Ooi HS, Ariyaratne PN, Vega VB, Luo Y, Tan PY, Choy PY, Wansa KDSA, Zhao B, Lim KS, Leow SC, Yow JS, Joseph R, Li H, Desai K V., Thomsen JS, Lee YK, Karuturi RKM, Herve T, Bourque G, Stunnenberg HG, Ruan X, Cacheux-Rataboul V, Sung WK, Liu ET, Wei CL, Cheung E, Ruan Y. 2009. An oestrogen-receptor-α-bound human chromatin interactome. Nature 462:58–64.

36. Mirny LA, Imakaev M, Abdennur N. 2019. Two major mechanisms of chromosome organization. Curr Opin Cell Biol 58:142–152.

37. Gunsalus LM, Keiser MJ, Pollard KS. 2023. In silico discovery of repetitive elements as key sequence determinants of 3D genome folding. Cell Genomics 3:100410.

38. Fudenberg G, Kelley DR, Pollard KS. 2020. Predicting 3D genome folding from DNA sequence with Akita. Nat Methods 17:1111–1117.

39. Schwessinger R, Gosden M, Downes D, Brown RC, Oudelaar AM, Telenius J, Teh YW, Lunter G, Hughes JR. 2020. DeepC: predicting 3D genome folding using megabase-scale transfer learning. Nat Methods 17:1118–1124.

40. Dixon JR, Selvaraj S, Yue F, Kim A, Li Y, Shen Y, Hu M, Liu JS, Ren B. 2012. Topological domains in mammalian genomes identified by analysis of chromatin interactions. Nature 485:376– 380.

41. Wolff J, Rabbani L, Gilsbach R, Richard G, Manke T, Backofen R, Grüning BA. 2020. Galaxy HiCExplorer 3: A web server for reproducible Hi-C, capture Hi-C and single-cell Hi-C data analysis, quality control and visualization. Nucleic Acids Res 48:W177–W184.

42. Durand NC, Shamim MS, Machol I, Rao SSP, Huntley MH, Lander ES, Aiden EL. 2016. Juicer Provides a One-Click System for Analyzing Loop-Resolution Hi-C Experiments. Cell Syst 3:95– 98.

43. Servant N, Varoquaux N, Lajoie BR, Viara E, Chen CJ, Vert JP, Heard E, Dekker J, Barillot E. 2015. HiC-Pro: An optimized and flexible pipeline for Hi-C data processing. Genome Biol 16:1– 11.

44. Zheng Y, Ay F, Keles S. 2019. Generative modeling of multi-mapping reads with mHi-C advances analysis of Hi-C studies. Elife 8:1–29.

45. Dossin FDM, Schenkman S. 2005. Actively Transcribing RNA Polymerase II Concentrates on Spliced Leader Genes in the Nucleus of *Trypanosoma cruzi*. Eukaryot Cell 4:1–11.

46. Peng T, Hou Y, Meng H, Cao Y, Wang X, Jia L, Chen Q, Zheng Y, Sun Y, Chen H, Li T, Li C. 2023. Mapping nucleolus-associated chromatin interactions using nucleolus Hi-C reveals pattern of heterochromatin interactions. Nat Commun 14:1–15.

47. Van Koningsbruggen S, Gierliński M, Schofield P, Martin D, Barton GJ, Ariyurek Y, Den Dunnen JT, Lamond AI. 2010. High-resolution whole-genome sequencing reveals that specific chromatin domains from most human chromosomes associate with nucleoli. Mol Biol Cell 21:3735–3748.

48. Carmo-Fonseca M, Mendes-Soares L, Campos I. 2000. To be or not to be in the nucleolus. Nat Cell Biol 2:E107–E112.

49. Pita S, Dıaz-Viraque F, Iraola G, Robello C. 2019. The tritryps comparative repeatome: Insights on repetitive element evolution in trypanosomatid pathogens. Genome Biol Evol 11:546–551.

50. Berná L, Chiribao ML, Greif G, Rodriguez M, Alvarez-Valin F, Robello C. 2017. Transcriptomic analysis reveals metabolic switches and surface remodeling as key processes for stage transition in *Trypanosoma cruzi*. PeerJ 2017:1–32.

51. Lima PLC de, Lopes L de S, Roson JN, Borges A, Bellini NK, Tahira A, Silva MS da, Pires D, Elias MC, Cunha JPC da. 2024. Comprehensive Analysis of Nascent Transcriptome Reveals Diverse Transcriptional Profiles Across the *Trypanosoma cruzi* Genome Underlining the Regulatory Role of Genome Organization, Chromatin Status, and Cis-Acting Elements. bioRxiiv.

52. Müller LSM, Cosentino RO, Förstner KU, Guizetti J, Wedel C, Kaplan N, Janzen CJ, Arampatzi P, Vogel J, Steinbiss S, Otto TD, Saliba AE, Sebra RP, Siegel TN. 2018. Genome organization and DNA accessibility control antigenic variation in trypanosomes. Nature 563:121–125.

53. Faria J, Luzak V, Müller LSM, Brink BG, Hutchinson S, Glover L, Horn D, Siegel TN. 2021. Spatial integration of transcription and splicing in a dedicated compartment sustains monogenic antigen expression in African trypanosomes. Nat Microbiol 6.

54. Nora EP, Lajoie BR, Schulz EG, Giorgetti L, Okamoto I, Servant N, Piolot T, Van Berkum NL, Meisig J, Sedat J, Gribnau J, Barillot E, Blüthgen N, Dekker J, Heard E. 2012. Spatial partitioning of the regulatory landscape of the X-inactivation centre. Nature 485:381–385.

55. Rowley MJ, Corces VG. 2018. Organizational principles of 3D genome architecture. Nat Rev Genet 19:789–800.

56. Ramírez F, Bhardwaj V, Arrigoni L, Lam KC, Grüning BA, Villaveces J, Habermann B, Akhtar A, Manke T. 2018. High-resolution TADs reveal DNA sequences underlying genome organization in flies. Nat Commun 9.

57. Szabo Q, Bantignies F, Cavalli G. 2019. Principles of genome folding into topologically associating domains. Sci Adv 5.

58. Rowley MJ, Nichols MH, Lyu X, Ando-Kuri M, Rivera ISM, Hermetz K, Wang P, Ruan Y, Corces VG. 2017. Evolutionarily Conserved Principles Predict 3D Chromatin Organization. Mol Cell 67:837–852.e7.

59. Ren L, Wang Y, Shi M, Wang X, Yang Z, Zhao Z. 2012. CTCF mediates the cell-type specific spatial organization of the kcnq5 locus and the local gene regulation. PLoS One 7:1–9.

60. Nichols MH, Corces VG. 2015. A CTCF Code for 3D Genome Architecture. Cell 162:703–705.

61. Taddei A, Gasser SM. 2012. Structure and function in the budding yeast nucleus. Genetics 192:107–129.

62. Yague-Sanz C, Migeot V, Larochelle M, Bachand F, Wéry M, Morillon A, Hermand D. 2023. Chromatin remodeling by Pol II primes efficient Pol III transcription. Nat Commun 14.

63. Thoru Pederson. 2011. The nucleolus. Nature 153:687–688.

64. Smircich P, Eastman G, Bispo S, Duhagon MA, Guerra-Slompo EP, Garat B, Goldenberg S, Munroe DJ, Dallagiovanna B, Holetz F, Sotelo-Silveira JR. 2015. Ribosome profiling reveals translation control as a key mechanism generating differential gene expression in *Trypanosoma cruzi*. BMC Genomics 16:1–14.

65. Bickmore WA. 2013. The spatial organization of the human genome. Annu Rev Genomics Hum Genet 14:67–84.

66. Cremer T, Cremer M. 2010. Chromosome territories. Cold Spring Harb Perspect Biol 2:1–22.

67. Bolger AM, Lohse M, Usadel B. 2014. Trimmomatic: A flexible trimmer for Illumina sequence data. Bioinformatics 30:2114–2120.

68. Knight PA, Ruiz D. 2013. A fast algorithm for matrix balancing. IMA J Numer Anal 33:1029– 1047.

69. Lowe TM, Chan PP. 2016. tRNAscan-SE On-line: integrating search and context for analysis of transfer RNA genes. Nucleic Acids Res 44:W54–W57.

70. Marck C, Kachouri-Lafond R, Lafontaine I, Westhof E, Dujon B, Grosjean H. 2006. The RNA polymerase III-dependent family of genes in hemiascomycetes: Comparative RNomics, decoding strategies, transcription and evolutionary implications. Nucleic Acids Res 34:1816–1835.

71. Wolff J, Backofen R, Grüning B. 2022. Loop detection using Hi-C data with HiCExplorer. Gigascience 11:1–9.

72. Simonis M, Klous P, Splinter E, Moshkin Y, Willemsen R, De Wit E, Van Steensel B, De Laat W. 2006. Nuclear organization of active and inactive chromatin domains uncovered by chromosome conformation capture-on-chip (4C). Nat Genet 38:1348–1354.

